# The landscape of actionable genomic alterations in cell-free circulating tumor DNA from 21,807 advanced cancer patients

**DOI:** 10.1101/233205

**Authors:** Oliver A. Zill, Kimberly C. Banks, Stephen R. Fairclough, Stefanie A. Mortimer, James V. Vowles, Reza Mokhtari, David R. Gandara, Philip C. Mack, Justin I. Odegaard, Rebecca J. Nagy, Arthur M. Baca, Helmy Eltoukhy, Darya I. Chudova, Richard B. Lanman, AmirAli Talasaz

## Abstract

Cell-free DNA (cfDNA) sequencing provides a non-invasive method for obtaining actionable genomic information to guide personalized cancer treatment, but the presence of multiple alterations in circulation related to treatment and tumor heterogeneity pose analytical challenges. We present the somatic mutation landscape of 70 cancer genes from cfDNA deep-sequencing analysis of 21,807 patients with treated, late-stage cancers across >50 cancer types. Patterns and prevalence of cfDNA alterations in major driver genes for non-small cell lung, breast, and colorectal cancer largely recapitulated those from tumor tissue sequencing compendia (TCGA and COSMIC), with the principle differences in alteration prevalence being due to patient treatment. This highly sensitive cfDNA sequencing assay revealed numerous subclonal tumor-derived alterations, expected as a result of clonal evolution, but leading to an apparent departure from mutual exclusivity in treatment-naïve tumors. To facilitate interpretation of this added complexity, we developed methods to identify cfDNA copy-number driver alterations and cfDNA clonality. Upon applying these methods, robust mutual exclusivity was observed among predicted truncal driver cfDNA alterations, in effect distinguishing tumor-initiating alterations from secondary alterations. Treatment-associated resistance, including both novel alterations and parallel evolution, was common in the cfDNA cohort and was enriched in patients with targetable driver alterations. Together these retrospective analyses of a large set of cfDNA deep-sequencing data reveal subclonal structures and emerging resistance in advanced solid tumors.

## Introduction

Genomic analysis of cell-free DNA (cfDNA) from advanced cancer patients allows the identification of actionable alterations shed into the circulation and may provide a global summary of tumor heterogeneity without an invasive biopsy (*1*). Plasma cfDNA analysis can provide insights from genomic information shed from multiple lesions within a patient, but this broader level of insight can introduce added complexity. Indeed, most clinical cfDNA sequencing is performed on patients with advanced or metastatic disease, often at the second or later line of therapy.

As a recently developed testing method, clinical cfDNA sequencing has repeatedly been benchmarked against tissue sequencing, but these performance comparisons are challenged by temporal and spatial heterogeneity in tumors (*2*–*6*). In addition, circulating tumor DNA (ctDNA) may be undetectable when shedding of tumor DNA is nominal, such as when therapy stabilizes tumor growth (*7, 8*). Recent efforts to globally characterize tumor heterogeneity using both plasma cfDNA analysis and multi-region tumor sequencing highlight the complementary nature of the two approaches (*9*–*11*). However, there is a paucity of cfDNA data sets large enough to evaluate the similarity of tumor-initiating alterations (“truncal drivers”) in solid tumor cancers to those found in the cfDNA of advanced cancer patients. Among the various methods available for cfDNA analysis, targeted panel deep-sequencing assays that utilize extensive error-correction methods provide the depth (sensitivity) and genomic breadth necessary to optimally survey tumor-derived genomic alterations in plasma cfDNA, even at low allelic fractions (*12*–*15*).

In order to elucidate the landscape of truncal driver mutations in cfDNA, we first evaluated the extent of detectable tumor heterogeneity in a large set of cfDNA deep-sequencing data, considering how varying ctDNA levels across patients might impact cfDNA variant detection. To distinguish truncal driver mutations from secondary resistance mutations, we developed methods to infer clonality and driver status of tumor mutations from cfDNA. We then examined the similarity of cfDNA patterns of common driver alterations in a large cohort of advanced, previously treated solid tumor patients to those found in treatment-naïve tumor tissue compendia. Finally, we explored the landscape of resistance to targeted therapies that was highly apparent in the large cfDNA cohort.

## Results

### Somatic genomic alterations in cfDNA across 21,807patients

Somatic cfDNA alterations were detected in 85% (18,503/21,807) of patients across all cancer types, ranging from 51% for glioblastoma to 93% for small cell lung cancer (**Figure 1A**). Half of the reported somatic cfDNA alterations had VAF <0.41% (range 0.03% - 97.6%; **Figure 1B**). Alteration-positive samples had on average three or four alterations detected (median=3; mean=4.3; range 1-166), including copy number amplifications (CNAs) (**Figure S1B**). Using the maximum somatic cfDNA VAF as an approximate measure of the level of ctDNA in a sample, we examined ctDNA level per indication. Although most of the major cancers (bladder, liver, prostate, gastric, NSCLC, melanoma, breast) had similar average levels of ctDNA, brain cancers had significantly lower levels (Wilcoxon p = 0.006 in comparison with renal) putatively owing to the blood-brain barrier whereas CRC and SCLC had significantly higher levels than all other indications examined (Wilcoxon p < 0.008 in comparison with bladder). (Similar patterns have been previously reported by (*40*), but without sufficient sample size to determine statistical significance.) This variation in ctDNA levels suggested the possibility of inter-indication variability in variant detection and therefore in ability to estimate tumor mutation load from cfDNA.

**Figure 1.**
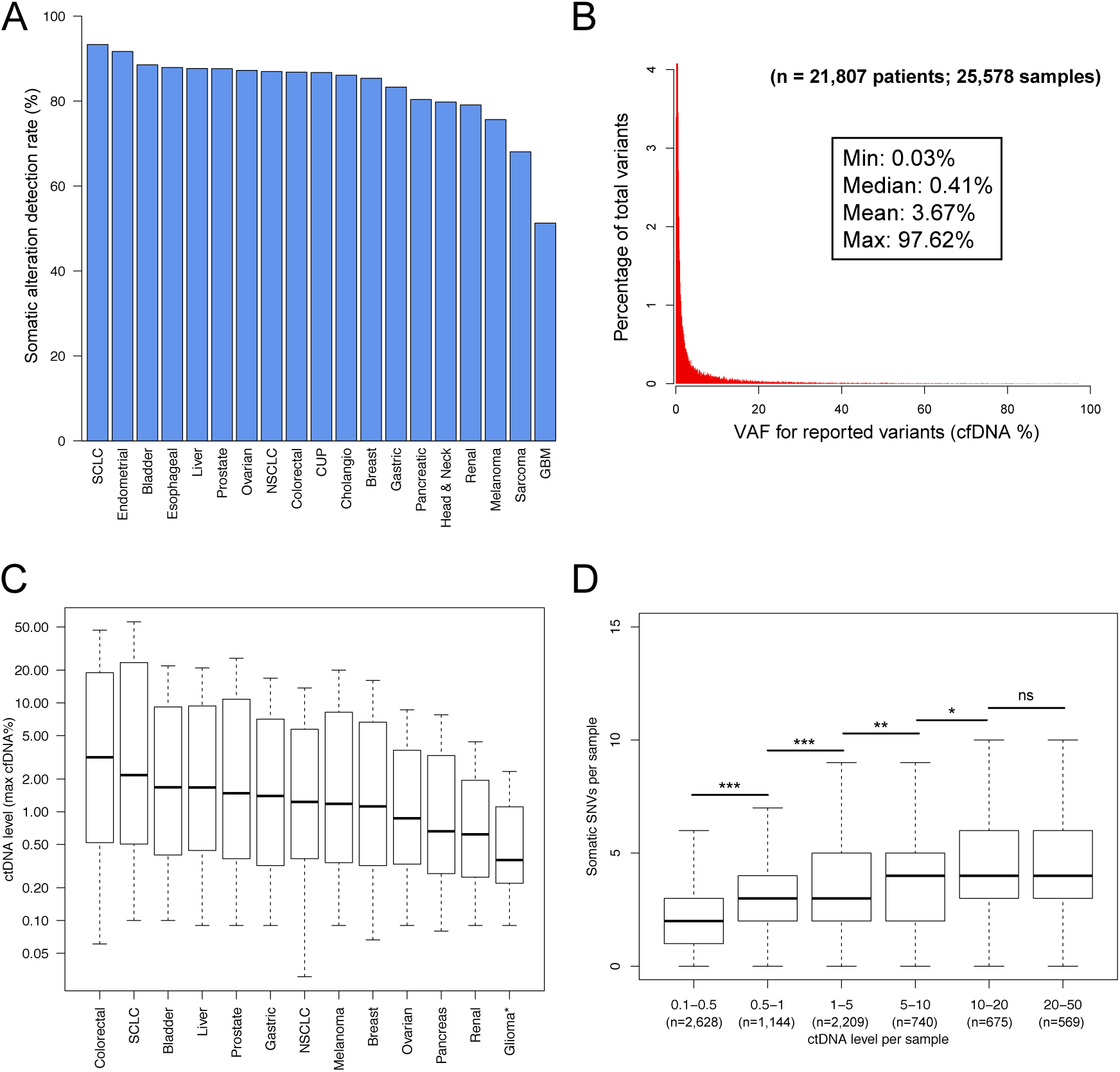
CfDNA alteration detection and ctDNA levels in 21,807 advanced-stage cancer patients. **(A)** Somatic cfDNA alteration detection rates per cancer type in the 21,807-patient cfDNA cohort. Percentages of alteration-positive samples are indicated. Note that the last 16,939 consecutive samples (Nov 2015-Sept 2016) were analyzed with version 2.9 of the cfDNA test, whereas the previous 8,639 samples were analyzed with earlier versions of the cfDNA test (see Table S3). SCLC, Small Cell Lung Cancer; CUP, Cancer of Unknown Primary; GBM, Glioblastoma. **(B)** VAF distribution for all somatic SNVs, indels, and fusions detected by the cfDNA test. **(C)** Distributions of ctDNA level per indication. CtDNA levels were significantly higher in colorectal cancer and SCLC and signficantly lower in glioma/GBM (“Glioma*”) than in the other cancers shown (Wilcoxon rank sum test). Sample numbers per indication are: Colorectal, 1,991; SCLC, 267; Bladder, 210; Liver, 210; Prostate, 909; Gastric, 260; NSCLC, 8,078; Melanoma, 410; Breast, 3,301; Ovarian, 594; Pancreas, 867; Renal, 220; Glioma/GBM, 107. **(D)** The number of somatic cfDNA SNVs detected per sample versus increasing levels of circulating tumor DNA (ctDNA). CfDNA NSCLC samples (n=8,132) are binned by their maximum VAF (cfDNA %) on the x-axis. Asterisks indicate significance levels from pairwise comparisons using the Wilcoxon rank sum test (“ns”, not significant).

Indeed, the number of cfDNA alterations per sample notably increased in samples with higher levels of ctDNA (**Figure 1D**), likely reflecting improved detection of genomic alterations when tumors shed more DNA into circulation. We tested whether ctDNA levels affected tumor mutation load estimates by considering the number of SNVs per sample in cfDNA NSCLC cases. When the mutations from TCGA NSCLC cases were filtered to those lying within the cfDNA panel regions (107 kb of sequence for reported variants), this cohort’s average mutation load was 18 mutations/Mb rather than the value of 9 mutations/Mb derived from whole-exome analysis. This increased average mutation load and compressed dynamic range across the cohort suggested a reduced accuracy for estimating average mutation load using these genomic regions, likely because they enrich for cancer driver mutations rather than passenger mutations. Nonetheless, the median tumor mutation load estimated from cfDNA steadily increased from 18.7 muts/Mb (2 SNVs per sample) in low-ctDNA NSCLC samples to 37.4 muts/Mb (4 SNVs per sample) in high-ctDNA samples (**Figure 1D and S1C**). As described below, at least part of the increase in mutation load in high-ctDNA samples was due to increased detection of subclonal variants (**Figure S2**).

CfDNA copy number analysis of 18 genes across four major cancer indications (lung, breast, colorectal, prostate) revealed amplification patterns consistent with known driver alterations in each indication (**Figure S3**). For example, *EGFR* was the most commonly amplified gene in lung cancers, *MYC* and *FGFR1* were the most commonly amplified gene in breast cancer, and *AR* was the most commonly amplified gene in prostate cancer. Notably, some established driver genes tended to have higher amplification levels than other genes that reflected indication-specific biology. For instance, *ERBB2* (HER2) had the highest average amplification levels in breast cancer and CRC but had middling amplification levels in lung and prostate cancers.

### Comparisons of alteration patterns across alteration types in cfDNA versus TCGA

To determine whether alteration patterns found in cfDNA recapitulated those found in published tissue sequencing studies, the frequencies of SNVs and indels in commonly mutated driver genes were compared to the frequencies found in TCGA. Highly similar mutation patterns were observed for *TP53* and *EGFR* (Pearson r=0.94 and r=0.78, respectively; **Figure 2A and B**), as well as for *KRAS, BRAF,* and *PIK3CA* (r=0.99, 0.99, and 0.94, respectively; data not shown). *EGFR*^T790M^ and *EGFR*^C797S^, treatment-induced resistance mutations, were more frequent in the heavily pre-treated cfDNA NSCLC cases (10%) relative to the untreated TCGA NSCLC cases (0.3%). Excluding T790M and C797S resistance alterations, the Pearson correlation for mutation frequencies in the tyrosine kinase domain of *EGFR* (exons 18-24) rose from 0.78 to 0.90.

**Figure 2.**
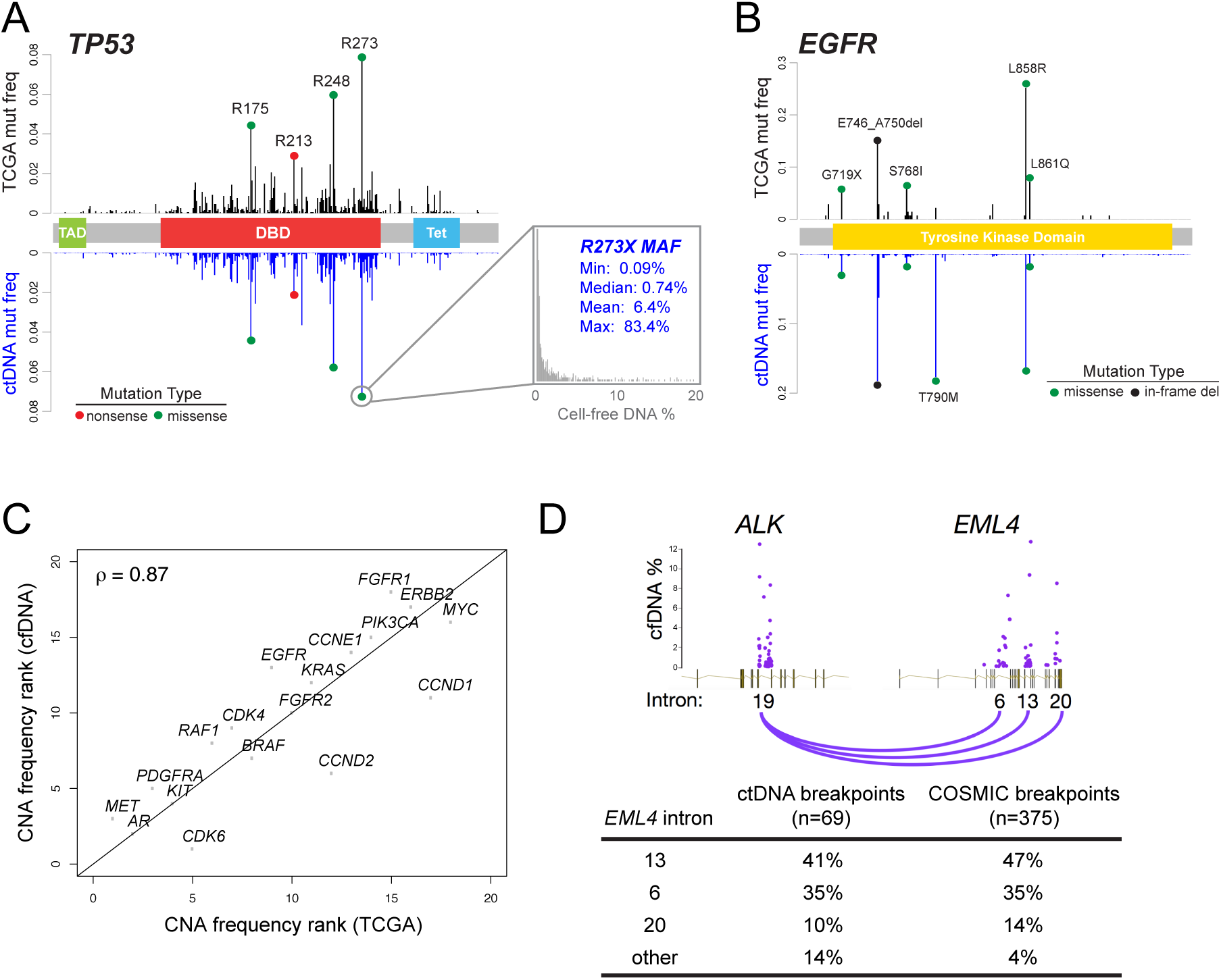
Comparison of cfDNA alteration patterns to tumor tissue alteration patterns in TCGA and COSMIC. **(A)** Per-codon mutation frequencies for SNVs in the *TP53* coding sequence [cfDNA n=14,696 SNVs (10,574 samples); tissue/TCGA n=1,951 SNVs (1,845 samples)]. **(B)** Per-codon mutation frequencies for the *EGFR* tyrosine kinase domain (exons 18-24) [cfDNA n=3,098 SNVs/indels (2,095 samples; tissue/TCGA n=112 SNVs/indels (96 samples)]. **(C)** Rank-by-rank comparison of amplification frequencies in breast cancer from the cfDNA cohort (1,010 patients with amplifications out of 2,808 patients) versus the tissue/TCGA cohort (413 samples with amplifications out of 816 profiled samples). **(D)** Comparison of *EML4-ALK* fusion breakpoints for cfDNA versus tissue (COSMIC). Top panel: schematic showing breakpoints versus VAF, expressed as cfDNA percentage, for *EML4-ALK* fusions detected in cfDNA. Bottom panel: breakpoint frequency per *EML4* intron; tissue data were compiled by COSMIC database (http://cancer.sanger.ac.uk/cosmic) from various literature sources.

Breast cancer often harbors actionable CNAs in *ERBB2* (HER2). We compared the ranks of amplification frequencies in breast cancer patients for the 18 CNA genes assayed by the cfDNA assay to the same genes in TCGA (amplification status determined by GISTIC), and found high rank correlation (ρ=0.86; **Figure 2C**). Similarly, 5-10% of lung adenocarcinoma (LUAD) is driven by targetable kinase gene fusions. To compare the patterns of gene fusions found in cfDNA to those found in tissue, we determined the frequencies per intron of breakpoints in the three most commonly observed fusions among lung cancer patients in the cfDNA cohort: *EML4-ALK*, *CCDC6*-*RET*, and *KIF5B*-*RET*. Breakpoint locations for all three fusions were strongly correlated with the frequencies of breakpoints found in published tissue data (r=0.98; **Figure 2D, Table S5**).

### CfDNA clonality and driver alteration prevalence in cfDNA versus TCGA

The abundance of advanced, treated cancer cases in the cfDNA cohort was expected to contribute additional subclonal variants when variant detection in cfDNA was not limited by low ctDNA levels. The trend toward increased numbers of cfDNA variants in high-ctDNA samples (**Figure 1D**) and the observation of frequent resistance alterations (**Figure 2B**) suggested that comparisons of this cfDNA cohort to large tumor tissue cohorts like TCGA should account for a higher level of mutational heterogeneity in the cfDNA cases. In order to compare the prevalences of common alterations between the large tissue cohorts of earlier stage tumors (TCGA) with those of the advanced-stage tumors in the cfDNA cohort, accounting for the potentially increased mutational heterogeneity in cfDNA, we derived a “cfDNA clonality” metric using the VAF/maximum VAF ratio that would allow us to infer the likely cancer-cell fraction of mutations present in the tumor (see Methods). We noted that mutated oncogenes such as *EGFR* could be subsequently amplified, which could inflate the cfDNA VAF leading to an inaccurate clonality estimate. Closer examination of the VAF/CN relationship revealed two separate nonlinear behaviors: non-linearity of amplified driver mutations at high VAF and high copy number, and clearly subclonal alterations with low VAF that occurred subsequent to, or in a separate subclone from, the amplification (**Figure 3A, Figures S6-S9**). Our model therefore takes into account these non-linearities by log-linear copy number normalization of the VAF for driver variants and by holding out variants that initially appear subclonal from the normalization procedure. We then examined the cfDNA clonality distributions for the most frequently mutated genes in lung adenocarcinoma (LUAD), breast cancer, and CRC to understand how this metric related to well-known biological properties among cancer mutations (**Figure S4**).

**Figure 3.**
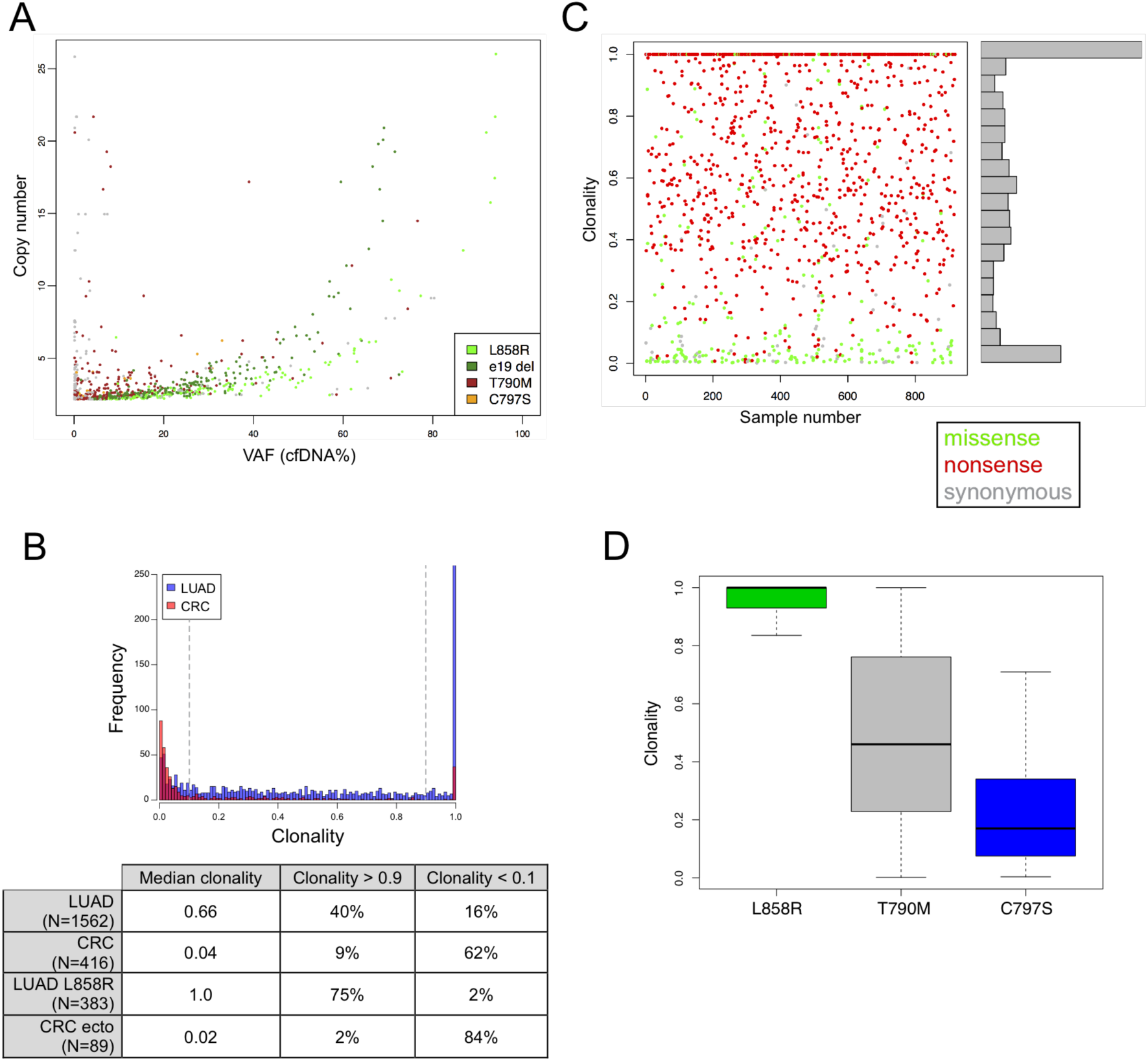
Determination of cfDNA clonality. **(A)** Copy number versus VAF for EGFR in NSCLC. Note the two different non-linear behaviors among low-VAF and high-VAF alterations. **(B)** CfDNA clonality is plotted for all *EGFR* mutations in LUAD (blue) or CRC (pink). Thresholds for clonality filtering are indicated by vertical grey lines. The median clonality and percentage of mutations that were clonal (clonality>0.9) or subclonal (clonality<0.1) for all *EGFR* mutations, for L858R alone in LUAD, or for recurrent ectodomain mutations (“ecto”) in CRC are shown in table below. **(C)** CfDNA clonality for all APC SNVs in CRC cases. Nonsense variants are colored red, missense variants are green, and synonymous are grey. **(D)** CfDNA clonality of a canonical *EGFR* driver alteration (L858R) and two resistance alterations (T790M, C797S) in cfDNA from 1,119 NSCLC samples. Note the relationship between variant clonality and order of treatment with a given therapy is consistent with the sequential emergence of each resistance alteration (erlotinib is given to patients with EGFR^L858R^; EGFR^T790M^ confers resistance to erlotinib, and patients with EGFR^L858R,T790M^ can then be given osimertinib; EGFR^C797S^ confers resistance to osimertinib).

*EGFR* mutations were among the most prevalent alteration across the cfDNA cohort, but had different expected cohort-level behaviors in LUAD versus CRC. In LUAD, *EGFR*-activating mutations should occur frequently as drivers and these mutations should tend to be clonal. In CRC, recurrent *EGFR* extracellular domain mutations would generally be expected to be acquired resistance alterations in patients treated with anti-EGFR antibodies such as cetuximab, and therefore should tend to be subclonal. As predicted, cfDNA *EGFR* alterations in LUAD were predominantly clonal, whereas in CRC they were predominantly subclonal (**Figure 3B**). Direct comparison of the clonality distributions of *EGFR*^L858R^ in LUAD versus *EGFR* ectodomain mutations in CRC showed an even more striking dichotomy (data not shown). In CRC, alterations in the common driver genes *APC*, *TP53*, and *KRAS* were predominantly clonal (**Figure S4**). Strikingly, nonsense mutations in *APC* had a strong tendency to be clonal (median clonality = 0.72), and had significantly higher average clonality than *APC* missense or synonymous alterations (median clonality = 0.07; Wilcoxon p < 10^-6^), further confirming that the cfDNA clonality metric reflected the expected behaviors of tumor-derived alterations (**Figure 3C**). In breast cancer, alterations in the common driver genes *PIK3CA*, *AKT1* and *TP53* showed strong tendencies toward clonality (**Figure S4**). Additionally, we compared the clonality distributions in LUAD of known *EGFR* driver (e19 del, L858R, etc.) and *EGFR* resistance (T790M, C797S) alterations. Again, as expected, the driver alterations showed a strong tendency toward clonality and the resistance alterations showed a strong tendency toward subclonality (**Figure 3D, Figure S4).**

Comparisons of mutation prevalence per gene between the cfDNA and TCGA cohorts, accounting for cfDNA clonality, revealed that the prevalences of most major driver alterations in NSCLC, breast cancer, and CRC were overall similar (Pearson correlation, r=0.85; **Figure S5, Table S6**). Some genes had significant differences in mutation prevalence between cohorts (Chi-square test), which largely reflected differences in patient demographics (i.e., prior treatment). Notable differences included *EGFR* and *KRAS* alterations in NSCLC (cfDNA: 43% *EGFR*-mutant, 16% *KRAS*-mutant; TCGA/tissue: 14% *EGFR*-mutant, 33% *KRAS*-mutant), a much higher frequency of *ESR1* mutations in cfDNA breast cancer samples (14%) than in TCGA samples (0.5%), and a substantially higher frequency of *TP53* mutations in cfDNA CRC samples.

### Mutual exclusivity analysis of driver alterations in cfDNA

To determine whether truncal driver alterations followed patterns of mutual exclusivity established in early-stage disease (i.e., TCGA studies), we performed mutual exclusivity analysis on common cfDNA alterations in LUAD, breast cancer, and CRC samples. Because driver status for CNAs is often unclear or ambiguously reported across studies, we developed and applied a cohort-level CNA driver identification method that retains statistical outliers relative to background aneuploidies to enrich the initial set of CNA calls for likely driver alterations (**Figure S10**, see Methods). Similarly, cfDNA SNVs, indels, and fusions were filtered to clonal alterations (clonality > 0.9) to enrich for likely truncal drivers (see Methods).

In LUAD, strong evidence for mutual exclusivity was observed in cfDNA across several pairs of genes (**Figure 4**). Importantly, the tendency for mutual exclusivity increased when comparing the post-clonality-filtering alterations to the pre-filtering alterations (**Figure S11 and S12, Tables S7-S12).** Of note, *KRAS* and *EGFR* were highly mutually exclusive in both cases, but with a 40x drop in the proportion of double-mutant [*KRAS-alt*; *EGFR-alt*] genotypes after filtering to clonal alterations. For *MET* and *EGFR*, a tendency toward alteration co-occurrence pre-filtering (FDR=6x10^-6^, logOR=0.6) was flipped to one of exclusivity after filtering (FDR=4x10^-3^, logOR=-1.3), suggesting that mutation co-occurrence before filtering was caused by subclonal resistance alterations, as opposed to co-occuring truncal mutations (the pre-filtered data had a high prevalence of *MET* amplifications and *EGFR*^T790M^, both of which are associated with resistance to erlotinib). A similar pattern of flipping from co-occurrence (non-significant) to exclusivity was seen for *ERBB2* and *EGFR* alterations (FDR=5x10^-7^, logOR=-2.6).

**Figure 4.**
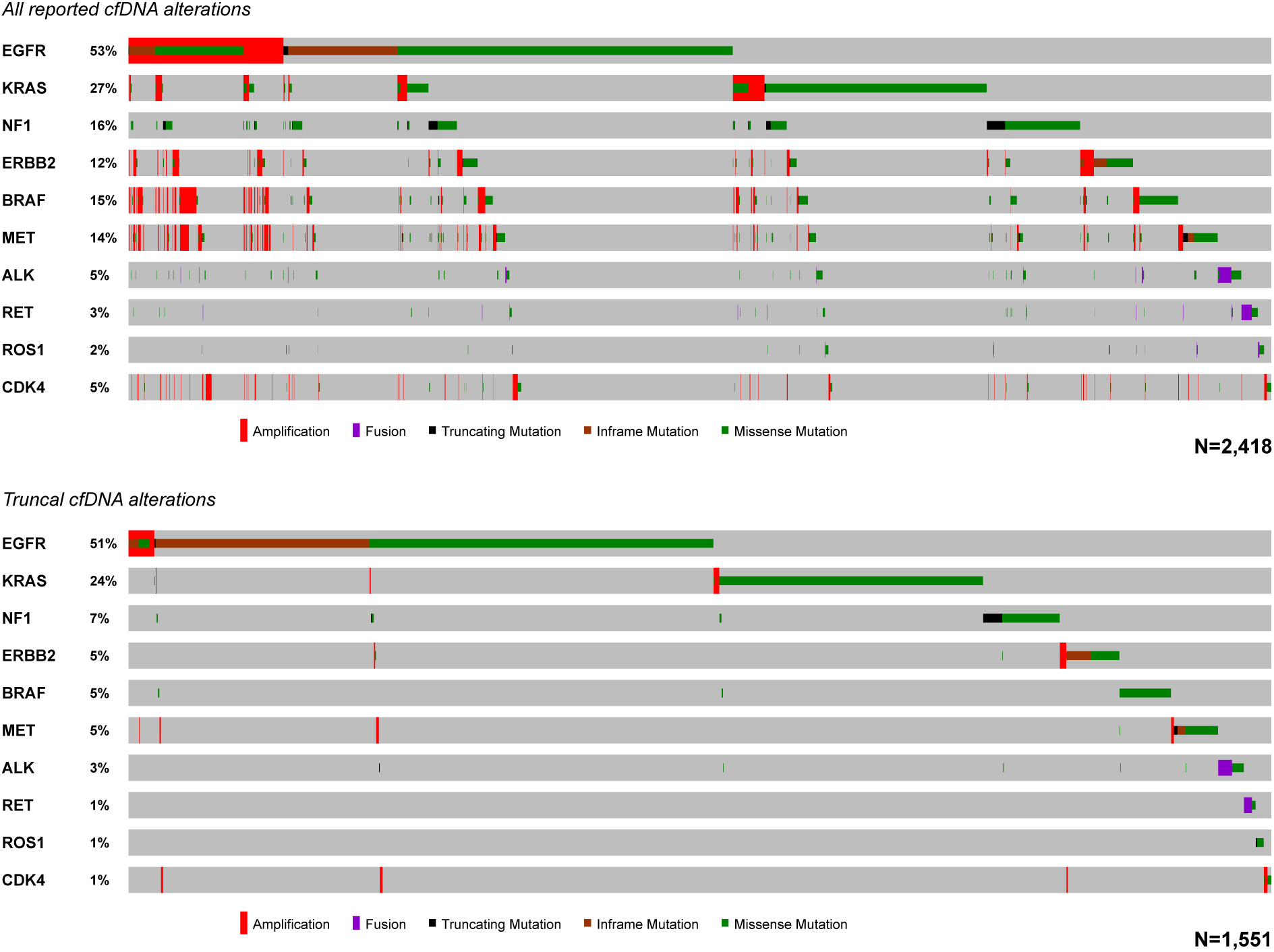
Mutual exclusivity analysis of cfDNA alterations for lung adenocarcinoma before and after filtering for clonality and CNA driver status. Numbers of patients are indicated at bottom right of each plot. The top oncoprint shows all alterations in alteration-positive patients, whereas the bottom oncoprint shows truncal SNVs, indels, and fusions (clonality > 0.9), and likely driver CNAs across patients that have at least one clonal driver alteration. Grey boxes indicate the absence of alterations, and the color/shape combinations corresponding to the various alteration types are indicated below each oncoprint. Frequencies of gene alterations within each plot are indicated at left (samples lacking clonal alterations in the selected genes were omitted).

In breast cancer, five driver genes (*ERBB2*, *FGFR1*, *BRCA1*, *BRCA2*, *AKT1*) showed tendencies toward mutual exclusivity with *PIK3CA* after clonality filtering. Exclusivity was not necessarily expected for *PIK3CA* alterations except with respect to *AKT1* mutations (*22*), reflecting the more complementary nature of driver alterations in this disease (**Figure S11 and S12, Tables S7-S12).** In CRC, *TP53*, *KRAS*, and *APC* alterations tended to co-occur, as expected (data not shown). Mutual exclusivity was observed between *KRAS* and *BRAF*, *ERBB2*, *NRAS*, and *MET*, similar to reports in tumor tissue. In the pre-filtered data, *KRAS* and *PIK3CA* tended to co-occur, but in the post-filtering data they showed a weak trend toward exclusivity, suggesting the presence of subclonal *KRAS* resistance alterations in the cfDNA CRC samples. We also noted that filtered *FGFR1* amplifications and *ERBB2* (HER2) amplifications showed a weak trend toward exclusivity in breast cancer (unadjusted p=0.04) and CRC, although in the latter case significance could not be readily assessed owing to the small number of certain genotype classes.

### The landscape of actionable resistance alterations in cfDNA

The large cfDNA cohort provided a unique opportunity to explore qualitatively and quantitatively the evolution of resistance alterations in patients who have progressed on targeted therapies in regular clinical practice. To estimate the frequency of actionable resistance alterations (defined as alterations that might influence a physician’s choice of therapy postprogression) in advanced, previously treated cancer patients, we identified known resistance alterations in the cfDNA cohort across six cancer types: NSCLC, breast cancer, CRC, prostate cancer, melanoma, and GIST. 3,397 samples of the 14,998 samples analyzed (22.6%) had at least one of the 134 known resistance alterations that were identified by the cfDNA test. The proportion of samples harboring likely resistance alterations increased when each indication was limited to samples harboring driver alterations with associated FDA-approved targeted drugs (hereafter, “on-label targetable driver alterations”), consistent with the resistance alterations having arisen due to therapy (**Figure 5A and B, Table S13**). The most common resistance alterations were *EGFR*^T790M^ and *MET* CNA in NSCLC, *AR* ligand-binding-domain SNVs in prostate cancer, *KRAS*^G12/G13/Q61^ in CRC, and *ESR1*^L536/Y537/D538^ in breast cancer (**Figure 5B, Table S13).**

**Figure 5.**
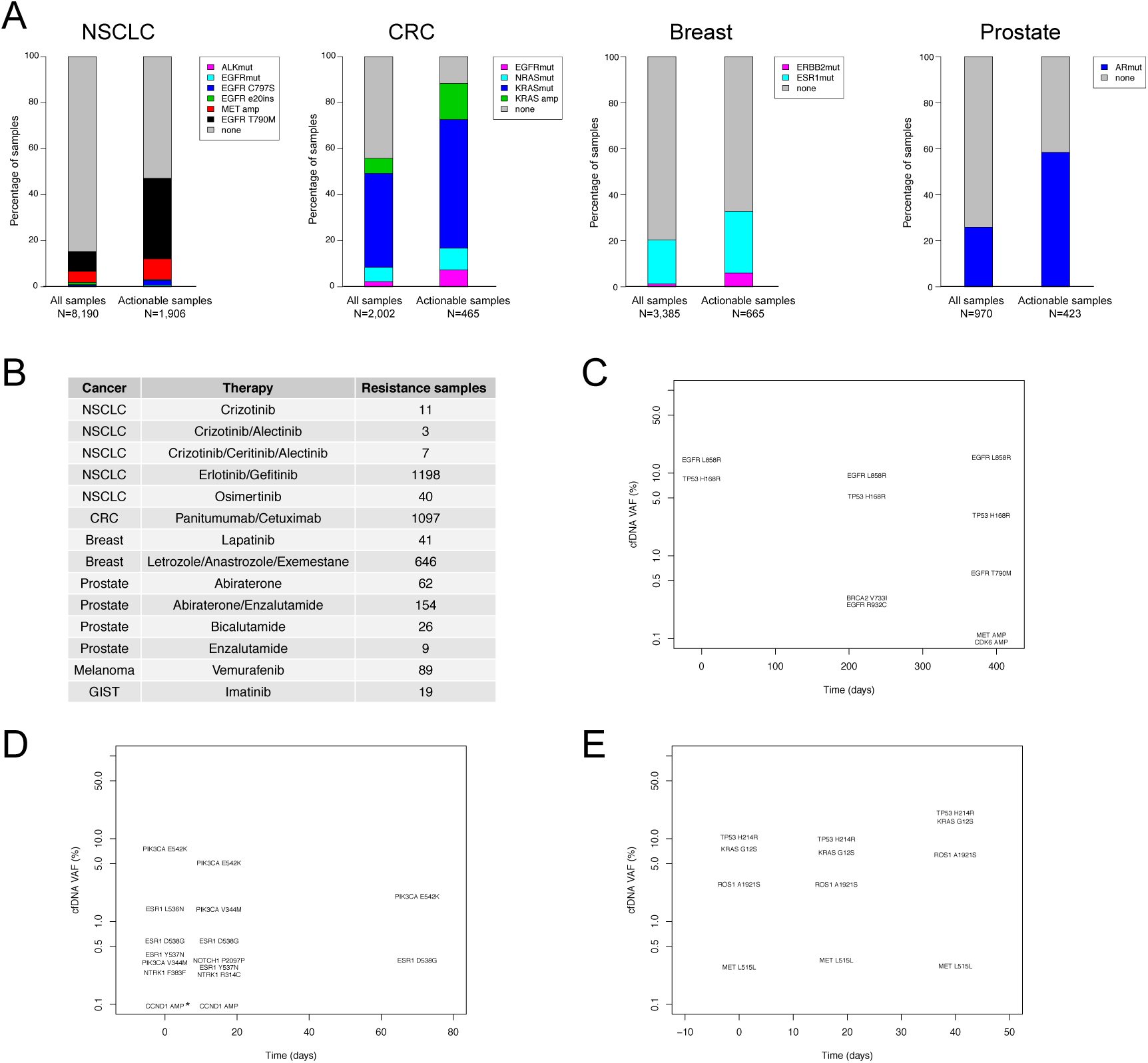
CfDNA landscape of resistance to on-label therapies across cancer types. **(A)** Landscape of resistance alterations in cfDNA. Numbers of patients with various resistance alterations (y-axis, left) to targeted therapies (x-axis, bottom) in six common cancer types (top) are plotted. Note that some patients harbored multiple distinct resistance alterations. Cancer type/genotype categories are: *AR*-mutant prostate cancer, *ALK*-fusion-positive NSCLC, *EGFR*-mutant NSCLC, breast cancer with *ESR1* or *ERBB2* (HER2) mutations, CRC, *BRAF*-mutant melanoma, and *KIT*-mutant gastrointestinal stromal tumor (GIST). The “EGFRmut” category for NSCLC includes variants A722V, L747P, L747S, V769M, T854A, T854S (18 mutations in total). **(B)** The numbers of samples harboring putative resistance mutations found in cfDNA and the corresponding on-label targeted therapies across six cancer types. **(C)** Longitudinal monitoring analysis of a patient with emerging resistance (T790M) in the third draw after presumptive EGFR inhibitor therapy indicated by L858R. Note that y-axis is log scale. **(D)** Monitoring analysis showing polyclonal *ESR1* mutations, which confer resistance to aromatase inhibitors, and possible patient response to therapy over time. Asterisk indicates four additional amplifications were detected in the first sample. **(E)** Monitoring analysis showing stability of the clonal stucture (rela tive VAF) over consecutive draws.

Although some resistance mutations can either occur as primary, truncal drivers or emerge secondarily upon treatment (e.g., *KRAS* ^G12X/G13X^ in CRC can be a truncal driver or emerge upon treatment with cetuximab), the cfDNA clonality metric helped distinguish resistance alterations whose emergence was likely caused by therapy pressure (**Figure S13**). A conservative estimate, focusing on clearly subclonal SNVs (clonality <0.1), was that at least 18.6% (ranging 10-34% across cancer types) of samples with on-label targetable alterations (381/2,053) had emerging secondary resistance alterations to those on-label therapies. Further, 24% of those resistance-harboring samples (91/381) had >1 alteration associated with resistance to the same therapy, suggesting independent evolution in distinct tumor lesions (*23*) or sequential treatment with distinct therapies targeted to the same gene. For example, one NSCLC patient had an *EML4-ALK* fusion (VAF of 7.1%) and *ALK* SNVs reported to confer resistance to crizotinib (L1196M, 2.5%), crizotinib/alectinib (I1171T, 0.1%), and crizotinib/ceritinib/alectinib (G1202R, 5%). In another example, the treatment history of certain patients harboring *EGFR*^L858R^ or *EGFR*^e19del^ driver alterations was immediately apparent by the combined presence of secondary *EGFR*^T790M^ and tertiary *EGFR*^C797S^ resistance alterations (24 patients had both *EGFR*^T790M^ and *EGFR*^C797S^ – 21 patients had these two variants in *cis*, the other 3 were in *trans*). The cfDNA clonality of *EGFR*^C797S^ was generally lower in those cases than that of *EGFR*^T790M^ (**Figure 3C**), consistent with tumor evolution following sequential lines of treatment with erlotinib/afatinib/gefitinib, followed by osimertinib at progression.

Novel resistance alterations were also identified in this clinical cohort, including: *ERBB2*^T798I^ (analogous to *EGFR*^T790M^) which causes resistance to an ERBB2 tyrosine kinase inhibitor; *MET*^D1228N^, *MET*^Y1230H^, and *MET*^G1163R^ (analogous to *ALK*^G1202R^ and *ROS1*^G2302R^) causing resistance of *MET* exon 14-mutated NSCLC to a next-generation MET inhibitor; and five *FGFR2* mutations (V564F, N549H, K641R, E565A, and L617V) shown to drive resistance to a selective pan-FGFR inhibitor (*24*–*26*); and the recurrent *EGFR* ectodomain mutations V441D/G, which arise in the setting of cetuximab resistance in CRC but are not yet characterized as functional (*27*). These putative resistance alterations were consistently subclonal relative to the original driver alteration and many were missed by single-metastatic-site tissue biopsy but confirmed by repeat biopsy or biopsy of multiple metastases at autopsy (*24, 25, 28).*

To illustrate the temporal dimension of the cfDNA landscape, we identified patients with multiple tests and significant clonal structure apparent in their ctDNA. These longitudinal cases illustrated emerging or polyclonal resistance after presumptive targeted therapy (**Figures 5C and D**), as well as stability of VAF estimates and clonality estimates over time (**Figure 5E**).

## Discussion

Much of our understanding of cancer genomes is derived from early-stage, treatment-naïve cancers via consortia efforts such as TCGA. However, the desire to increase treatment efficacy in advanced cancers that likely have evolved considerably from baseline has led to a recent shift to “real world” cancer genomics studies focused on the realities of the clinic, yet grounded in lessons from earlier-stage cancers (*29, 30*). It is becoming increasingly clear that obtaining comprehensive genomic assessments, across heterogeneous tumor subclones, will be necessary for tailoring effective therapies for advanced cancer patients (*9, 10*).

We have provided the largest cohort-level snapshot of genomic alterations in advanced cancer patients by cfDNA analysis in real-life clinical practice. Our results demonstrate that patterns and frequencies of truncal driver alterations in advanced cancers reflect patterns found in early-stage disease, but also reflect the increased complexity of advanced, treated cancers. We found that cfDNA alterations (SNVs and small, activating indels) in *TP53*, *EGFR*, *KRAS*, *PIK3CA*, *BRAF* strongly correlated with TCGA tissue alterations (r=0.90-0.99, **Figure 2A and B**), and that correlations for amplification frequency ranks in breast cancer and locations of intronic fusion breakpoints in NSCLC were similarly high (**Figure 2C and D**). The high sensitivity of the cfDNA assay combined with the more evolved advanced cancers tested at progression, which have greater numbers of mutations than earlier-stage, treatment-naïve cancers, may explain most of the differences in estimated tumor mutation load versus TCGA (*10, 30*). Importantly, we show that accurate estimation of tumor mutation load from plasma cfDNA will require taking ctDNA level into account, as the two factors are correlated (**Figure 1D**). Our estimates of ctDNA levels are based on the copy-number-adjusted allelic frequency of cfDNA somatic alterations, and future studies should also consider allele-specific molecule counts (germline allele imbalance) in estimates of tumor DNA in circulation.

Our inference of tumor mutation clonality based on copy-number-adjusted relative cfDNA VAF (**Figure 3**) enabled a recapitulation of mutual exclusivity among truncal driver mutations and facilitated identification of subclonal emerging resistance alterations (**Figure 4, Figure 5C, Figures S6-S8**). These results suggest that mutation clonality, as it exists in tumor tissue, can be inferred from analysis of relative cfDNA VAFs, as has been previously hypothesized (*31*). High accuracy in VAF estimation is likely key to the success of this approach, and notably, VAFs measured by the cfDNA NGS assay used in this study show good agreement with digital droplet PCR (*32, 33*). This approach points to the possibility of analyzing the clonal structures of tumors from cfDNA sequencing data, unencumbered by the complications of tumor heterogeneity and tumor impurity introduced by single-region tissue sampling. Future studies of under-explored biological factors, such as the variability of cfDNA shedding across patients and the uniformity of cfDNA shedding across distinct tumor sites harboring genetically distinct clones, could enable statistical modeling of tumor clonal structures using cfDNA VAFs and cfDNA molecule counts per locus or per allele.

The most notable differences in prevalence of driver alterations between cfDNA and tissue cohorts were *EGFR* and *KRAS* alterations in LUAD (whose prevalences were flipped), and the higher frequency of *ESR1* mutations in cfDNA breast cancer samples and of *TP53* mutations in the cfDNA CRC samples. The higher *EGFR* alteration prevalence in LUAD cfDNA was likely due to a population bias resulting from clinicians ordering the cfDNA test at progression on an EGFR TKI (median time between diagnosis and plasma collection of 335 days). This is supported by *EGFR*^T790M^ being the one of the most common *EGFR* variants in the cfDNA cohort, second only to *EGFR* exon 19 deletion driver mutations. Screening known *EGFR*-mutant NSCLC patients at progression for resistance mutations is routine practice whereas re-profiling *KRAS*-mutant NSCLC patients would generally not be done, leading to an over-representation of *EGFR* driver mutations, and the concomitant under-representation of *KRAS* mutations, in this cohort. Similarly, the higher frequency of *ESR1* mutations (a documented resistance mechanisms to aromatase inhibitors) likely reflects the clinical application of ctDNA assays at progression. There are several possible explanations for the higher *TP53* prevalence in cfDNA CRC samples relative to TCGA: stage III/IV tumors, which predominate the cfDNA cohort, may have higher frequencies of *TP53* alterations than Stage I/II tumors; more subclonal *TP53* mutations may have been detected due to the high sensitivity of the cfDNA assay; or some *TP53* mutations may stem from somatic myeloid malignancies known as clonal hematopoiesis of indeterminate potential (*34, 35*). The most likely explanation is that the TCGA cohort (< 300 samples) underestimates *TP53* mutation prevalence in CRC relative to the larger 1,374 sample GENIE cohort, the latter reporting a 68.5% mutation prevalence for this gene (*36*).

The prevalence of actionable resistance alterations that could be linked to FDA-approved therapies or clinical trials of novel targeted agents is a key, and clinically important, finding of this cfDNA study. Nearly one in four cfDNA-alteration-positive patients (22.6%) across six cancer indications had one or more resistance alterations to an FDA-approved on-label therapy (**Figure 5A and B**), which would increase options for clinical decision-making. The significant enrichments for these candidate resistance alterations when cohorts were subset to patients with actionable driver alterations suggested that they were indeed linked to prior patient treatment. Our estimates of the frequencies of secondary, rather than primary, resistance alterations (10-34% of patients across the six cancer types) were likely conservative, as we examined only low-level subclonal SNVs (clonality < 0.1), in part because accurate assessment of clonality for CNAs remains difficult. As expected, the prevalence of resistance alterations was higher in cfDNA than TCGA/tissue, as these alterations would not be present in early stage tissue biopsies. For instance, the high frequency of cfDNA *ESR1* mutations in breast cancer patients likely reflected prior treatment with aromatase inhibitors. Additionally, *EGFR*^T790M^ was one of the most common *EGFR* mutations found in the cfDNA NSCLC cohort (8% of patients), but was seen in only two patients from the TCGA tissue NSCLC cohort (0.3%).

Although our cohort of clinically ordered cfDNA tests is uncontrolled, its large size provides a realistic cross-section of patients with advanced disease at the forefront of cancer care. Interpretation of our findings should take into consideration the selection biases related to cfDNA test ordering patterns in clinical practice and other potential limitations. Prevalences of genomic alterations may be biased by preferential ordering for patients with certain demographic characteristics, such as non-smoking females with NSCLC (thereby enriching for *EGFR* mutation over *KRAS* mutation). Plasma-based genotyping is often ordered at progression in advanced cancer, and thus is biased toward higher prevalences of resistance alterations, as discussed above. Although the cfDNA panel (70 genes) was focused primarily on the actionable portion of the cancer genome, a tradeoff of its relatively small size may be somewhat reduced power and accuracy to infer mutation clonality relative to an assay with a larger genomic footprint. However, it is currently impractical to perform whole-exome sequencing of cfDNA at >15,000X coverage.

This clinical cfDNA cohort represents the largest sequencing landscape of resistance in advanced cancer patients, and builds upon the body of primary driver alterations characterized by the TCGA, GENIE, and other projects. As such, a portion of this database has been included in the Blood Profiling Atlas in Cancer, a National Cancer Moonshot Initiative (*37*). Improved detection of resistance mutations may facilitate enrollment in clinical trials and enable the development of more accurate biomarkers of response to therapy (*24, 38, 39*). Therefore cfDNA, or other minimally invasive techniques, address a real and unmet need, since it is essential to provide real-time resampling of tumor genotype at the time of progression to guide subsequent therapeutic strategies.

## Materials and Methods

### Characteristics of the cfDNA cohort

A summary of the cfDNA cohort used in this study is provided in **Table 1**. The cohort was assembled from 21,807 consecutive cancer patients (25,578 total samples) tested on the Guardant Health cfDNA sequencing platform as part of clinical care. As such, this was an observational, non-interventional study. Patients consented to their test results being used in research, and all patient data were de-identified as per an institutional review board-approved protocol (Quorum Review IRB Protocol 30-001: Research Use of De-Identified Specimens and Data). Tests were ordered by 3,283 oncologists across the United States, Europe, Asia, and the Middle East from June 2014 to September 2016 (data freeze at 9/22/2016). Disease stage was confirmed for each patient by the provider to be advanced disease (Stage III/IV). Treatment histories and survival information were generally not available. Over 50 solid tumor types were represented [non-small cell lung cancer (NSCLC, 38%), breast cancer (18%), colorectal cancer (CRC, 11%), other (32%)], although many cancer types were represented by only a small number of samples. See **Tables S3 and S4** for comparisons of the cfDNA cohort characeristics with those of the pan-cancer TCGA cohort.

**Table 1.**
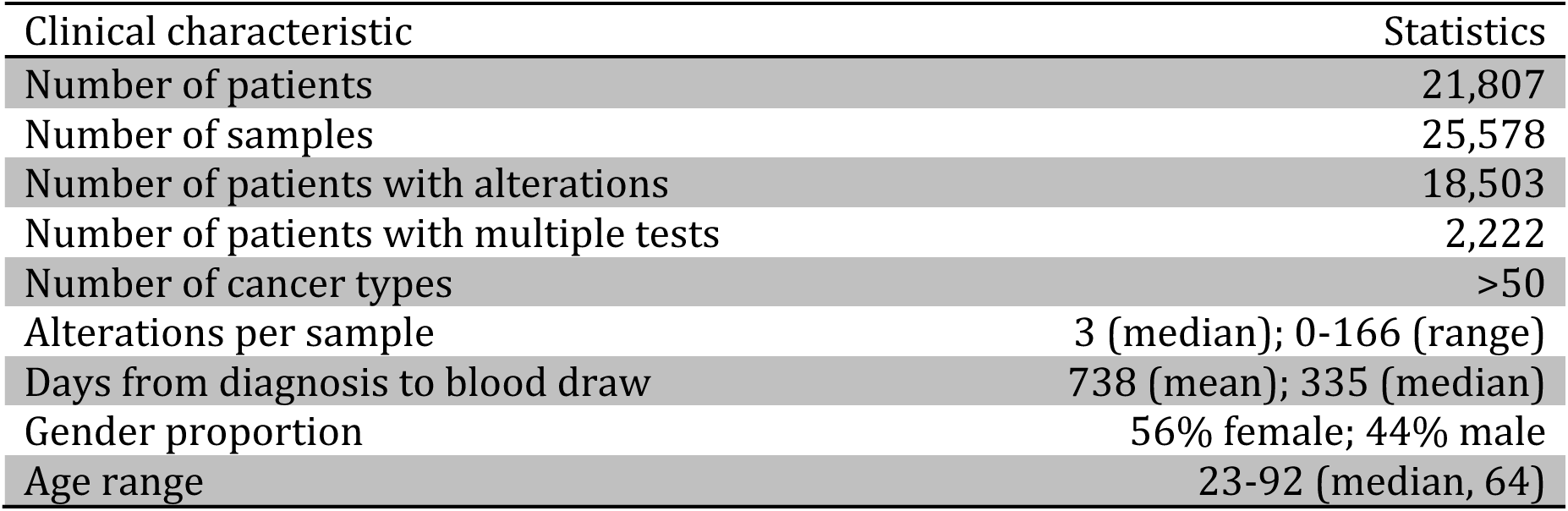
**Summary of the cfDNA cohort used in this study.** No treatment, follow-up, or outcome data were available.

### cfDNA sequencing assay and cfDNA variant calling

The clinical cfDNA sequencing platform (Guardant360) used in this study has been previously described (*13*). All analysis was performed in a Clinical Laboratory Improvement Amendments (CLIA)-certified, College of American Pathologists (CAP)-accredited, New York State Department of Health-approved clinical laboratory (Guardant Health, Inc., Redwood City, California). CfDNA was isolated from plasma after separating the plasma from 10mL whole blood per patient sample. Target capture of a 150kb region (all exons or critical exons of 70 genes, see **Tables S1 and S2**) was performed after minimal PCR amplification to ensure assay robustness and uniformity across samples. Samples were sequenced to an average depth of 15,000x raw read coverage per base pair. Somatic cfDNA alterations [single nucleotide variants (SNVs), indels, fusions, and copy number amplifications (CNAs)] were identified by a proprietary bioinformatics pipeline that reconstructs the original double-stranded cfDNA molecules present in a plasma sample, thereby transforming next-generation sequencing (NGS) read information into accurate, molecule-based variant calls. High specificity was achieved by combining molecular barcoding technology (in-line adapters are ligated immediately after cfDNA isolation, prior to PCR and target capture steps) and bioinformatics filtering of sequencing errors via statistical filtering of sequencing errors per-base-pair. Small variant detection was performed by comparing read-level and molecule-level characteristics to position-specific reference error noise profiles determined independently for each panel position sequencing training data from healthy donors. Observed positional error profiles were used to define calling cut-offs for variant detection. This process reduces the variant-detection error rate by several orders of magnitude (*16, 17, 13).* High sensitivity [80% probability of detecting SNVs at 0.3% variant allele fraction (VAF)] was achieved in part via end-to-end process improvements to ensure maximal recovery of cfDNA fragments throughout the assay.

To detect CNAs, probe-level unique molecule coverage was normalized for probe efficiency, GC content, and signal saturation, and the probe-level signals were then combined to report gene-level copy number. CNA calls were based on decision thresholds established from the training data, and considered both the absolute copy number deviation and the background variation within each sample’s own unique-molecule-counts distributions (diploid baseline). We note that the cfDNA bioinformatics pipeline determines unique molecules per base and determines VAF using mutant molecule counts (rather than read counts). This enables per-base per-patient detection-limit controls such that variants are only called if the number of unique mutant molecules is ≥2 and if the total unique-molecule count at a given genomic position are sufficient. Samples that have lower molecule diversity (unique molecules per base) have inherently lower sensitivity for variant calling, regardless of sequence-read depth.

### Comparison of cfDNA alterations to TCGA alterations

Genomic alterations in cfDNA were compared to those from primarily early-stage, treatment-naive tumors in TCGA studies (*18*). TCGA alterations from 9,077 samples across 27 cancer types were downloaded from the cBio portal (www.cbioportal.org) on 4/13/2016 using the cgdsr R package, and were filtered to the cfDNA panel footprint (Table S1) prior to analysis. For gene fusion comparisons, fusion breakpoints compiled from the COSMIC database (v76) were used rather than TCGA because of larger sample size (*19*).

Genes used for alteration prevalence comparisons were chosen based on the common driver genes in each cancer type documented by TCGA. Genes used for the mutual exclusivity analysis were chosen based on the overlap of commonly altered driver genes per cancer type, as shown by TCGA, and those alterations that are included in the cfDNA test. Oncoprints were generated using the cBio portal OncoPrinter web app (www.cbioportal.org/oncoprinter.jsp). Detection of short indels in tumor suppressor genes was not added to the cfDNA sequencing assay until November 2016 (Guardant360 v2.9, see **Table S1**), and therefore were not included in this study.

### Clonality inference of tumor-derived somatic alterations from cfDNA

The cfDNA clonality for somatic SNVs, indels, and fusions was initially defined as: ***alteration VAF / maximum somatic VAF in sample.*** In a patient with a copy-normal tumor (2n ploidy) with maximal cfDNA shedding from the tumor (such that non-tumor-derived DNA in circulation is negligible), clonal, heterozygous somatic variants should have cfDNA VAF equal to 50%. Assuming approximately proportional shedding across subclones and equivalent decay across cfDNA molecules in circulation, tumor-derived somatic variants should have cfDNA VAF proportional to their cancer-cell fraction (CCF) in the tumor. However, cancer genomes have frequent copy number alterations, which could impact the observed variant allele frequency in circulation. Point mutations that occur prior to an allele being amplified would have an inflated VAF in cfDNA relative to their tumor clonality, whereas mutations occurring after the amplification, or in a separate clone, would have VAF that remains in proportion to their clonality (assuming no additional copy number changes occur). Therefore, normalizing VAF by copy number only makes sense if a copy number alteration follows the point mutation.

Additionally, we noted that at high copy number the relationship between VAF and copy number becomes non-linear for amplified driver mutations (**Figure 3A**). Therefore a log transformation or non-linear regression approach is more appropriate than simple *VAF/CN* normalization for amplified mutations. We developed a cfDNA clonality model that considered both the relative timing of point mutation and amplification, and the non-linearity of *VAF/CN* at high copy number. Given the bifurcating VAF distribution for commonly amplified oncogenes (**Figure 3A, Figures S6-S9**), we normalized VAF to CN only if the initial VAF/maximum VAF ratio was >0.1, using a log-linear *VAF*/*ln(CN)* normalization for these mutations. Dividing each adjusted cfDNA alteration VAF by the adjusted maximum somatic VAF in a given sample then yielded the inferred alteration clonality present in the tumor. Note that we examined the frequency with which amplifications adversely impacted cfDNA clonality estimation by inflating the VAF, and found it to occur in <1% of samples (data not shown).

To determine reasonable thresholds for clonality filtering of driver alterations, the cfDNA clonality distributions were analyzed for SNVs in genes that were commonly mutated in CRC, NSCLC, and breast cancer, including either as primary drivers or as secondary resistance alterations (**Figure 3A and S4**). For most genes, the SNV clonality distributions were clearly bimodal, with the two maxima typically falling above 0.9 and below 0.1, respectively (**Figure 3A, Figure S4**). For the purposes of this study, alterations were considered clonal if their cfDNA clonality was >0.9 and subclonal if their cfDNA clonality was <0.1. Hence for the driver prevalence comparisons (**Figure S5, Table S6**), cfDNA alterations with clonality <0.1 were filtered out. For the mutual exclusivity analysis, “truncal cfDNA alterations” correspond to alterations with cfDNA clonality >0.9. Note that cfDNA clonality filtering does not uniformly remove alterations at low VAF, as low-VAF alterations remain in samples with low ctDNA. Since cfDNA mutation signatures in low-ctDNA samples are predominated by clonal tumor alterations, cfDNA clonality filtering effectively balances out the previous overrepresentation of subclonal, lower-VAF variants in high-ctDNA samples (**Figure S3).** We note that our method for estimating cfDNA clonality was independently developed and first described in multiple cancer types by (*20*), and cfDNA clonality determination has also recently been described in NSCLC by (*21*).

### Identification of CNA drivers in cfDNA

The cfDNA assay identifies CNAs based on statistical deviation of the normalized number of cfDNA molecules corresponding to a given gene above the diploid baseline. Whereas the CNA frequency comparison (**Figure 2C**) included all reported amplifications, for the mutual exclusivity analysis (**Figure 4**), cfDNA CNA calls were first filtered to likely driver alterations to exclude passenger alterations associated with aneuploidy. We used an algorthmic approach (“ctDriver”) that iteratively searched through CNAs per sample for outliers above an aneuploidy threshold determined from the ensemble of copy number calls in the cfDNA cohort (**Figure S10**). If the maximum CNA was >30% above the second-maximum CNA and >30% above the median copy number of all other CNAs in the sample, it was considered a driver. If the second-maximum was also >30% above the median copy number, both the maximum and second-maximum CNA were considered drivers. For example, if a sample had one CNA with copy number of 3.0 and five CNAs with copy number of 2.2, the former was considered focal and a likely driver alteration, whereas the other CNAs were considered likely to be caused by aneuploidy and were removed from the analysis. Additionally, any single CNA with cfDNA copy number above 5 (the 93^rd^ percentile of the overall copy number distribution) was considered a likely driver. We confirmed that these represented high-level outliers by fitting linear models of copy-number values that included the per-sample ctDNA estimate as an explanatory variable per cancer type. The result of this process narrowed the number of samples down to 1,396/6,159 samples with initial CNA calls (22%), and 1,396/16,185 initial CNA calls (9%) were deemed “driver CNAs” for the purposes of mutual exclusivity analysis.

### Resistance alterations and longitudinal analysis

Known drug resistance annotations (based on the literature) were tabulated for the 13,330 patients (11,539 with cancer alterations detected) with one of six cancer types with literature-documented resistance alterations and robust presence in the cfDNA cohort: NSCLC, breast cancer, CRC, prostate cancer, melanoma, gastrointestinal stromal tumor (GIST). Patients with at least one annotated resistance alteration were further examined for variant clonality (as above), presence of additional variants conferring resistance to the same therapy, and presence of additional variants conferring resistance to different therapies.

Of the 2,222 patients with multiple cfDNA tests, 731 had three or more tests (**Figure S1A**). Those 731 patients were then filtered for known major driver alterations in lung, colorectal, and breast cancer where tests were performed at relatively even time intervals. CfDNA alterations were then plotted for each test for patients representative of emerging resistance, stable clonal structure, and polyclonal resistance.

### Statistical analysis

Pearson correlations for SNVs/indels or fusion breakpoints, and Spearman correlations for amplification frequencies were calculated from tables of alteration frequencies in the cfDNA cohort versus in TCGA. Proportions for driver prevalence were compared using the Chi-square test. Mutation count, ctDNA level, and clonality distributions were compared using the Wilcoxon rank sum test, as described throughout the text. Fisher’s exact test was used to determine the statistical significance and log odds ratios for the mutual exclusivity analysis (**Figure 4, Tables S7-S12**), with adjustment for multiple comparisons within each cancer type using the Benjamini-Hochberg method (“false discovery rate”). All analyses were conducted in R.

## Acknowledgments

**General:** We thank all our colleagues at Guardant Health for their support, inspiration, and helpful discussions. We thank the patients, and the physicians who submitted samples that were deidentified and used in this analysis. The results published here are in part based upon data generated by the TCGA Research Network: http://cancergenome.nih.gov/.

## Author Contributions

O.Z., D.C., R.L., A.T. conceived and designed the study. O.Z., K.B., S.F., S.M., J.V., A.B., R.M., J.O., R.N., C.L., C.J., S.F., D.C. acquired, analyzed, or interpreted data, or made key methodological contributions. O.Z., K.B., D.C., R.L. wrote the paper. H.E., P.M., D.G., D.C., R.L., A.T. supervised the study.

## References

1. R. Lebofsky et al., Circulating tumor DNA as a non-invasive substitute to metastasis biopsy for tumor genotyping and personalized medicine in a prospective trial across all tumor types. Mol. Oncol. 9, 783–790 (2015).

2. M. Gerlinger et al., Intratumor heterogeneity and branched evolution revealed by multiregion sequencing. N. Engl. J. Med. 366, 883–892 (2012).

3. R. Govindan, Cancer. Attack of the clones. Science. 346, 169–170 (2014).

4. G. Gremel et al., Distinct subclonal tumour responses to therapy revealed by circulating cell-free DNA. Ann. Oncol. Off. J. Eur. Soc. Med. Oncol. 27, 1959–1965 (2016).

5. M. Murtaza et al., Multifocal clonal evolution characterized using circulating tumour DNA in a case of metastatic breast cancer. Nat. Commun. 6, 8760 (2015).

6. A. G. Sacher et al., Prospective Validation of Rapid Plasma Genotyping for the Detection of EGFR and KRAS Mutations in Advanced Lung Cancer. JAMA Oncol. 2, 1014–1022 (2016).

7. A. Vallée et al., Rapid clearance of circulating tumor DNA during treatment with AZD9291 of a lung cancer patient presenting the resistance EGFR T790M mutation. Lung Cancer. 91, 73–74 (2016).

8. A. Marchetti et al., Early Prediction of Response to Tyrosine Kinase Inhibitors by Quantification of EGFR Mutations in Plasma of NSCLC Patients: J. Thorac. Oncol. 10, 1437–1443 (2015).

9. C. Abbosh et al., Phylogenetic ctDNA analysis depicts early-stage lung cancer evolution. Nature. 545, 446–451 (2017).

10. M. Jamal-Hanjani et al., Tracking the Evolution of Non-Small-Cell Lung Cancer. N. Engl. J. Med. 376, 2109–2121 (2017).

11. E. Pectasides et al., Genomic Heterogeneity as a Barrier to Precision Medicine in Gastroesophageal Adenocarcinoma. Cancer Discov. (2017), doi:10.1158/2159-8290.CD-17-0395.

12. S. T. Kim et al., Prospective blinded study of somatic mutation detection in cell-free DNA utilizing a targeted 54-gene next generation sequencing panel in metastatic solid tumor patients. Oncotarget. 6, 40360–40369 (2015).

13. R. B. Lanman et al., Analytical and Clinical Validation of a Digital Sequencing Panel for Quantitative, Highly Accurate Evaluation of Cell-Free Circulating Tumor DNA. PloS One. 10, e0140712 (2015).

14. A. M. Newman et al., Integrated digital error suppression for improved detection of circulating tumor DNA. Nat. Biotechnol. 34, 547–555 (2016).

15. J. C. Thompson et al., Detection of Therapeutically Targetable Driver and Resistance Mutations in Lung Cancer Patients by Next-Generation Sequencing of Cell-Free Circulating Tumor DNA. Clin. Cancer Res. Off. J. Am. Assoc. Cancer Res. 22, 5772–5782 (2016).

16. I. Kinde, J. Wu, N. Papadopoulos, K. W. Kinzler, B. Vogelstein, Detection and quantification of rare mutations with massively parallel sequencing. Proc. Natl. Acad. Sci. U. S. A. 108, 9530–9535 (2011).

17. S. R. Kennedy et al., Detecting ultralow-frequency mutations by Duplex Sequencing. Nat. Protoc. 9, 2586–2606 (2014).

18. Cancer Genome Atlas Research Network, Comprehensive molecular profiling of lung adenocarcinoma. Nature. 511, 543–550 (2014).

19. S. A. Forbes et al., COSMIC: exploring the world’s knowledge of somatic mutations in human cancer. Nucleic Acids Res. 43, D805–8111 (2015).

20. O. A. Zill et al., Somatic genomic landscape of over 15,000 patients with advanced-stage cancer from clinical next-generation sequencing analysis of circulating tumor DNA. J. Clin. Oncol. 34 (2016) (available at http://meetinglibrary.asco.org/content/171265-176).

21. C. M. Blakely et al., Evolution and clinical impact of co-occurring genetic alterations in advanced-stage EGFR-mutant lung cancers. Nat. Genet. (2017), doi:10.1038/ng.3990.

22. Cancer Genome Atlas Network, Comprehensive molecular portraits of human breast tumours. Nature. 490, 61–70 (2012).

23. S. Misale, F. Di Nicolantonio, A. Sartore-Bianchi, S. Siena, A. Bardelli, Resistance to anti-EGFR therapy in colorectal cancer: from heterogeneity to convergent evolution. Cancer Discov. 4, 1269–1280 (2014).

24. L. Goyal et al., Polyclonal Secondary FGFR2 Mutations Drive Acquired Resistance to FGFR Inhibition in Patients with FGFR2 Fusion-Positive Cholangiocarcinoma. Cancer Discov. 7, 1–12 (2017). T798I

25. A. B. Hanker et al., An Acquired *HER2*^T798I^ Gatekeeper Mutation Induces Resistance to Neratinib in a Patient with HER2 Mutant?Driven Breast Cancer. Cancer Discov. 7, 575–585 (2017).

26. X. Lu et al., MET exon 14 mutation encodes an actionable therapeutic target in lung adenocarcinoma. Cancer Res. (2017), doi:10.1158/0008-5472.CAN-16-1944.

27. J. H. Strickler et al., Genomic landscape of cell-free DNA in patients with colorectal cancer. Cancer Discov. (2017), doi:10.1158/2159-8290.CD-17-1009.

28. M. Russo et al., Tumor Heterogeneity and Lesion-Specific Response to Targeted Therapy in Colorectal Cancer. Cancer Discov. 6, 147–153 (2016).

29. B. Vogelstein et al., Cancer genome landscapes. Science. 339, 1546–1558 (2013).

30. L. R. Yates et al., Genomic Evolution of Breast Cancer Metastasis and Relapse. Cancer Cell. 32, 169–184.e7 (2017).

31. Z. Piotrowska et al., Heterogeneity Underlies the Emergence of EGFRT790 Wild-Type Clones Following Treatment of T790M-Positive Cancers with a Third-Generation EGFR Inhibitor. Cancer Discov. 5, 713–722 (2015).

32. D. S. Hong et al., Phase IB Study of Vemurafenib in Combination with Irinotecan and Cetuximab in Patients with Metastatic Colorectal Cancer with BRAFV600E Mutation. Cancer Discov. 6, 1352–1365 (2016).

33. B. C. Ulrich et al., Cross-platform detection and quantification of actionable mutations in cell-free DNA shows high concordance and correlation between next-generation sequencing and droplet digital PCR. Cancer Res. 77, 5692 (2017).

34. D. P. Steensma et al., Clonal hematopoiesis of indeterminate potential and its distinction from myelodysplastic syndromes. Blood. 126, 9–16 (2015).

35. B. M. Zhang et al., IDH2 Mutation in a Patient with Metastatic Colon Cancer. N. Engl. J. Med. 376, 1991–1992 (2017).

36. AACR Project GENIE Consortium, AACR Project GENIE: Powering Precision Medicine through an International Consortium. Cancer Discov. (2017), doi:10.1158/2159-8290.CD-17-0151.

37. R. L. Grossman et al., Collaborating to Compete: Blood Profiling Atlas in Cancer (BloodPAC) Consortium. Clin. Pharmacol. Ther. (2017), doi:10.1002/cpt.666.

38. J. F. Gainor et al., Molecular Mechanisms of Resistance to First- and Second-Generation ALK Inhibitors in ALK-Rearranged Lung Cancer. Cancer Discov. 6, 1118–1133 (2016).

39. T. S. Mok et al., Osimertinib or Platinum-Pemetrexed in EGFR T790M-Positive Lung Cancer. N. Engl. J. Med. (2016), doi:10.1056/NEJMoa1612674.

40. C. Bettegowda et al., Detection of circulating tumor DNA in early- and late-stage human malignancies. Sci. Transl. Med. 6, 224ra24 (2014).

